# Cotton fabrics functionalized with hydroxyl-rich graphene derivatives and silver nanowires: washing resistance and preliminary antibacterial activity against *Escherichia coli*

**DOI:** 10.64898/2026.03.16.710677

**Authors:** A. H. Lima, D.B.R. Silva, G. R. Carvalho, A. C. P. Fernandes, C. T. Tavares, N. C. Vicentini, C. C. S. Cunha, R. A. Dias, A. Del’Duca, D. E. Cesar, A. S. A. Watanabe, W. G. Quirino

## Abstract

Cotton-based antimicrobial textiles are attractive for applications requiring improved microbiological control, but their performance depends on effective surface functionalization and retention of the active materials after use and washing. In this work, cotton fabrics were functionalized with hydroxyl-rich graphene oxide (HGO), hydroxyl-rich reduced graphene oxide (H-rGO), and silver nanowires (AgNWs), either individually or in combined treatments, to investigate their deposition onto the textile surface, washing resistance, and preliminary antibacterial activity. The treated fabrics were prepared by immersion-based coating procedures, and the persistence of the deposited materials after repeated washing was evaluated by UV-Vis analysis of the residual wash solutions. Surface morphology before and after washing was examined by scanning electron microscopy. The results showed that graphene-based coatings, particularly HGO, exhibited stronger retention on cotton fibers, while AgNWs were partially retained after repeated washing cycles. SEM images confirmed the deposition of AgNWs on the cotton surface and showed that part of the coating remained associated with the fibers after washing. A preliminary antibacterial assay against *Escherichia coli* indicated that nanomaterial-treated fabrics inhibited bacterial growth relative to untreated controls, with the combined HGO/AgNWs treatment showing the most promising inhibitory trend under the tested conditions. These findings demonstrate the feasibility of producing cotton fabrics functionalized with hydroxyl-rich graphene derivatives and silver nanowires, supporting their potential as proof-of-concept antibacterial textiles with partial washing resistance.

## 1. Introduction

Functionalized textiles with antimicrobial properties have attracted increasing attention due to their potential use in applications that require improved microbiological control, including healthcare-related materials, protective clothing, and high-contact reusable fabrics [1–3]. In this context, cotton is a particularly attractive substrate because it is inexpensive, widely available, biocompatible, and easy to process. However, as a hydrophilic fibrous material, cotton can also favor microbial retention and growth, which motivates the development of surface modification strategies capable of imparting antimicrobial activity without compromising the practical advantages of the textile substrate.

Among the nanomaterials investigated for antimicrobial surface engineering, graphene oxide (GO) and reduced graphene oxide (rGO) have emerged as promising candidates [2,4–10]. Their antimicrobial effects have been associated with direct physical interactions with microbial cells, including membrane disruption by sheet edges, as well as oxidative-stress-related mechanisms that depend on surface chemistry. In the present study, special attention was given to hydroxyl-rich graphene oxide (HGO) and hydroxyl-rich reduced graphene oxide (H-rGO), since the abundance of oxygen-containing groups may favor both interactions with the hydroxyl-rich cellulose surface and biologically relevant interfacial effects. The possibility of depositing these graphene-derived nanomaterials onto cotton fibers is particularly interesting for the preparation of washable functional textiles.

Silver-based nanostructures are also well known for their broad antimicrobial activity [3,11–13]. In particular, silver nanowires (AgNWs) combine the intrinsic antimicrobial properties of silver with a high-aspect-ratio morphology that may enhance surface coverage and contact with the treated substrate. Combining graphene-derived materials with AgNWs is therefore an attractive strategy for producing multifunctional coatings in which the graphene-based layer may improve interfacial adhesion and surface distribution, while silver contributes additional antibacterial activity.

Based on this rationale, the present work investigates the functionalization of cotton fabrics with HGO, H-rGO, and AgNWs, either individually or in combined treatments. The study focuses on three key aspects: material deposition onto the textile substrate, resistance of the treated fabrics to repeated washing, and preliminary antibacterial activity against Escherichia coli. By adopting this proof-of-concept approach, we aim to assess whether hydroxyl-rich graphene derivatives and silver nanowires can be incorporated into cotton fabrics in a way that preserves surface-associated material after washing while providing measurable antibacterial performance.

## 2. Materials and Methods

### 2.1 Synthesis of hydroxyl-rich graphene oxide, hydroxyl-rich reduced graphene oxide, and silver nanowires

Powder graphite, silver nitrate, sodium chloride, polyvinylpyrrolidone (PVP), ethylene glycol, acetone, and ethanol were used as received. Hydroxyl-rich graphene oxide (HGO) was prepared by a modified Hummers oxidation protocol, as previously reported by Lima et al. [14–16]. HGO dispersions were subsequently reduced with hydrazine hydrate to obtain stable aqueous dispersions of hydroxyl-rich reduced graphene oxide (H-rGO) [17,18]. Silver nanowires (AgNWs) were synthesized by a modified polyol route based on the procedure reported by Parente et al. [19]. The reaction was carried out in ethylene glycol at 160 °C under controlled stirring for 15 min, yielding AgNWs with an average length of approximately 10 μm. The dispersion was purified by repeated dilution in acetone, centrifugation at 2000 rpm for 5 min, and resuspension in deionized water for four cycles. The purified AgNWs were dispersed in ethanol and stored at 5 °C until use.

### 2.2 Preparation of treated cotton fabrics

Commercial cotton fabric (CF) was cut into 10 x 10 cm pieces and pre-washed with a 5% aqueous alkaline dextran detergent solution to remove contaminants from the fibers. The cleaned samples were dried at 50 °C overnight before surface treatment. For graphene-based functionalization, pre-washed CF samples were immersed in 50 mL aqueous suspensions of HGO (3 or 5 mg/mL) or H-rGO (0.1 mg/mL) for 30 min at room temperature. After immersion, the samples were removed from the bath and dried at 50 °C overnight. For silver nanowire deposition, pristine CF samples and graphene-treated samples were exposed to AgNWs dispersed in ethanol at 1 mg/mL. Before AgNWs treatment, the CF-HGO and CF-H-rGO samples were washed to remove excess nanosheets and dried again at 50 °C overnight. The samples were then immersed in 50 mL of AgNWs dispersion for 2 min, removed from the bath, and dried at 50 °C for 30 min. This deposition/drying cycle was repeated five times to improve AgNWs incorporation onto the textile surface, followed by a final washing step to remove excess nanowires.

### 2.3 Optical characterization and washing-resistance assay

UV-Vis spectroscopy measurements of HGO, H-rGO, and AgNWs dispersions were performed with an Ocean Optics USB2000 fiber spectrometer. For HGO and H-rGO measurements, 25 μL of each dispersion was diluted in 4 mL of deionized water, whereas 100 μL of AgNWs dispersion was diluted for analysis. To evaluate the resistance of the treated fabrics to washing, selected coated samples were subjected to five washing cycles. Each cycle corresponded to three repetitions of immersion in 80 mL of deionized water, vigorous vortex mixing for 1 min, and drying at 50 °C for 2 h. After each cycle, the residual wash water was collected and analyzed by UV-Vis spectroscopy.

### 2.4 Scanning electron microscopy

Scanning electron microscopy (SEM) was used to investigate the surface morphology of the cotton fabrics treated with HGO, H-rGO, AgNWs, and combined graphene-derivative/AgNWs coatings. SEM analyses were performed using a FEI Quanta 250 microscope operated at 30 kV. Prior to imaging, the samples were coated with a 40 nm Au film by sputtering.

### 2.5 Preliminary antibacterial assay against Escherichia coli

Bacterial isolates used in the microbiological assays were maintained in the Biology Techniques Laboratory at IF Sudeste MG. In the preliminary experiment retained in the present proof-of-concept manuscript, the growth of Escherichia coli on treated fabrics was compared among CF-HGO3, CF-AgNWs, and CF-HGO3/Ag. The control materials represented in the final graph were PP mask and washed cotton fabric (washed CF). Fabric samples were cut to 2 x 1 cm, sterilized by autoclaving after handling, and exposed to E. coli inoculum prepared from 0.5 McFarland bacterial suspension in BHI broth. The inoculum was applied by spraying 0.1 g of sterile BHI inoculum per 1 cm^2^ of fabric. The samples were placed on solid Mueller-Hinton agar in Petri dishes, incubated at 36.5 °C for 24 h, and then fixed with 2% paraformaldehyde. After collection, each sample was subjected to three sonication cycles with 1 min intervals, centrifuged for 5 min at 500 g, filtered through 0.2 μm polycarbonate membranes, stained with DAPI, and examined by epifluorescence microscopy. The biological comparison presented here is intentionally restricted to the first E. coli experiment to preserve a focused and self-consistent proof-of-concept narrative.

## 3. Results and Discussion

### 3.1 Optical characterization of the nanomaterials

The optical responses of HGO, H-rGO, and AgNWs were first evaluated to confirm the characteristic spectral features of the as-prepared materials. HGO exhibited the expected absorption bands centered around 230 and 300 nm, whereas the reduced material showed a red shift of the main absorption band to approximately 268 nm together with increased absorption over the visible range. This behavior is consistent with partial restoration of the conjugated carbon network after chemical reduction. The AgNWs dispersion displayed the characteristic plasmonic bands centered at approximately 350 and 378 nm, confirming the formation of elongated silver nanostructures with low contribution from side products.

**Figure 1.**
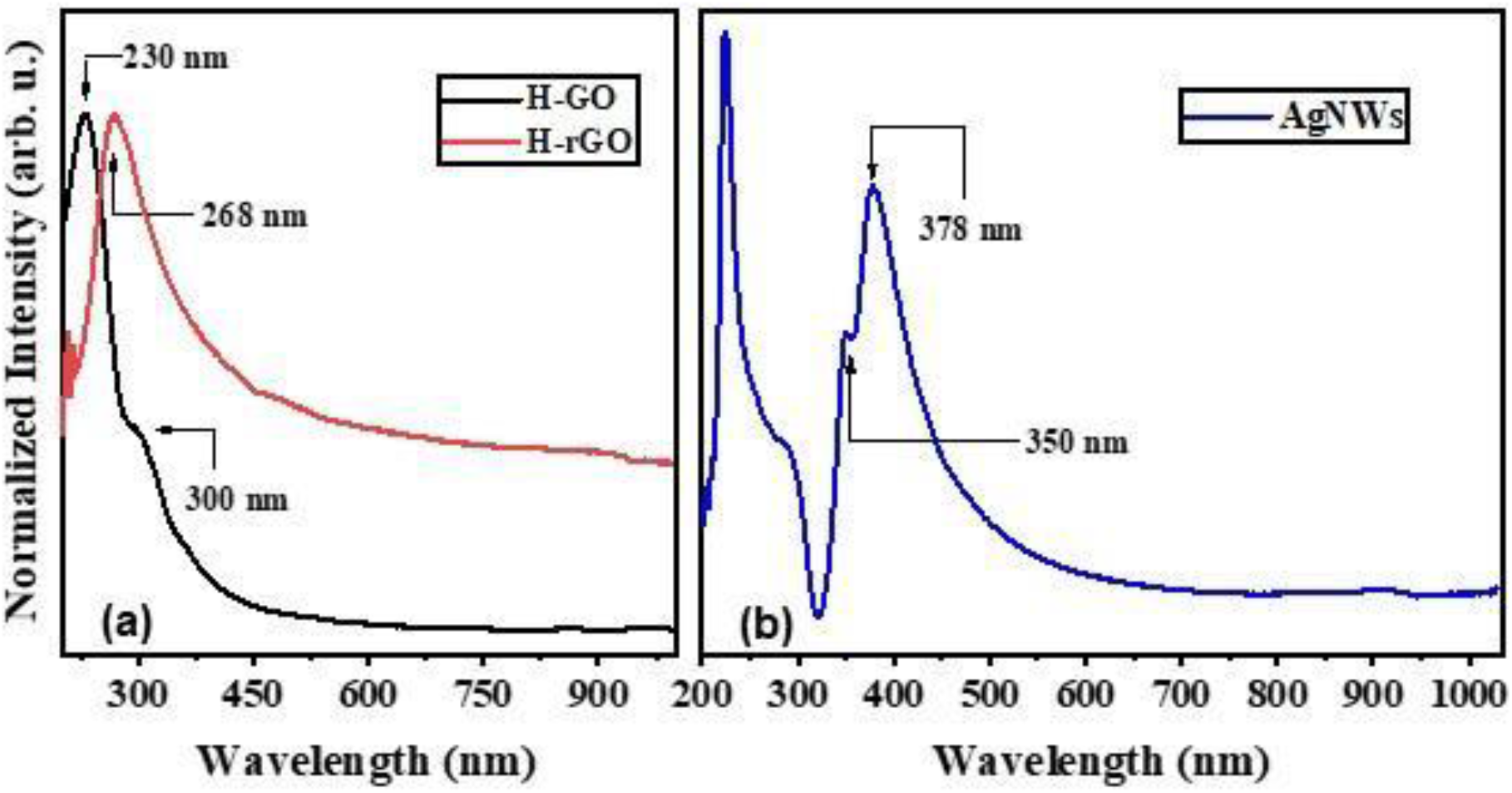
Ultraviolet-visible (UV-Vis) absorption spectra of the as-prepared nanomaterials: (a) hydroxyl-rich graphene oxide (HGO) and hydroxyl-rich reduced graphene oxide (H-rGO) dispersions and (b) raw silver nanowire (AgNWs) dispersion. Hydroxyl-rich graphene oxide (HGO) exhibits characteristic bands near 230 and 300 nm, hydroxyl-rich reduced graphene oxide (H-rGO) shows a red-shifted main band near 268 nm, and silver nanowires (AgNWs) exhibit plasmonic features centered at approximately 350 and 378 nm.

### 3.2 Washing resistance of treated cotton fabrics

The adhesion of the nanomaterials to cotton fibers was then investigated by monitoring the UV-Vis spectra of the residual washing water after successive washing cycles. For HGO-coated cotton, most of the loosely bound nanosheets were removed during the first washing cycle, whereas the following spectra became similar to those of washed untreated cotton. This pattern suggests that the remaining HGO nanosheets were more strongly associated with the cellulose surface than the excess material initially deposited. For the combined CF-HGO3/AgNWs sample, the residual wash water predominantly exhibited the characteristic plasmonic bands of AgNWs, indicating that silver nanowires were the main species released during subsequent washing. Together, these data indicate that graphene-based coatings, particularly HGO, showed stronger retention on cotton fibers than AgNWs alone, while silver nanowires remained partially retained after repeated washing.

**Figure 2.**
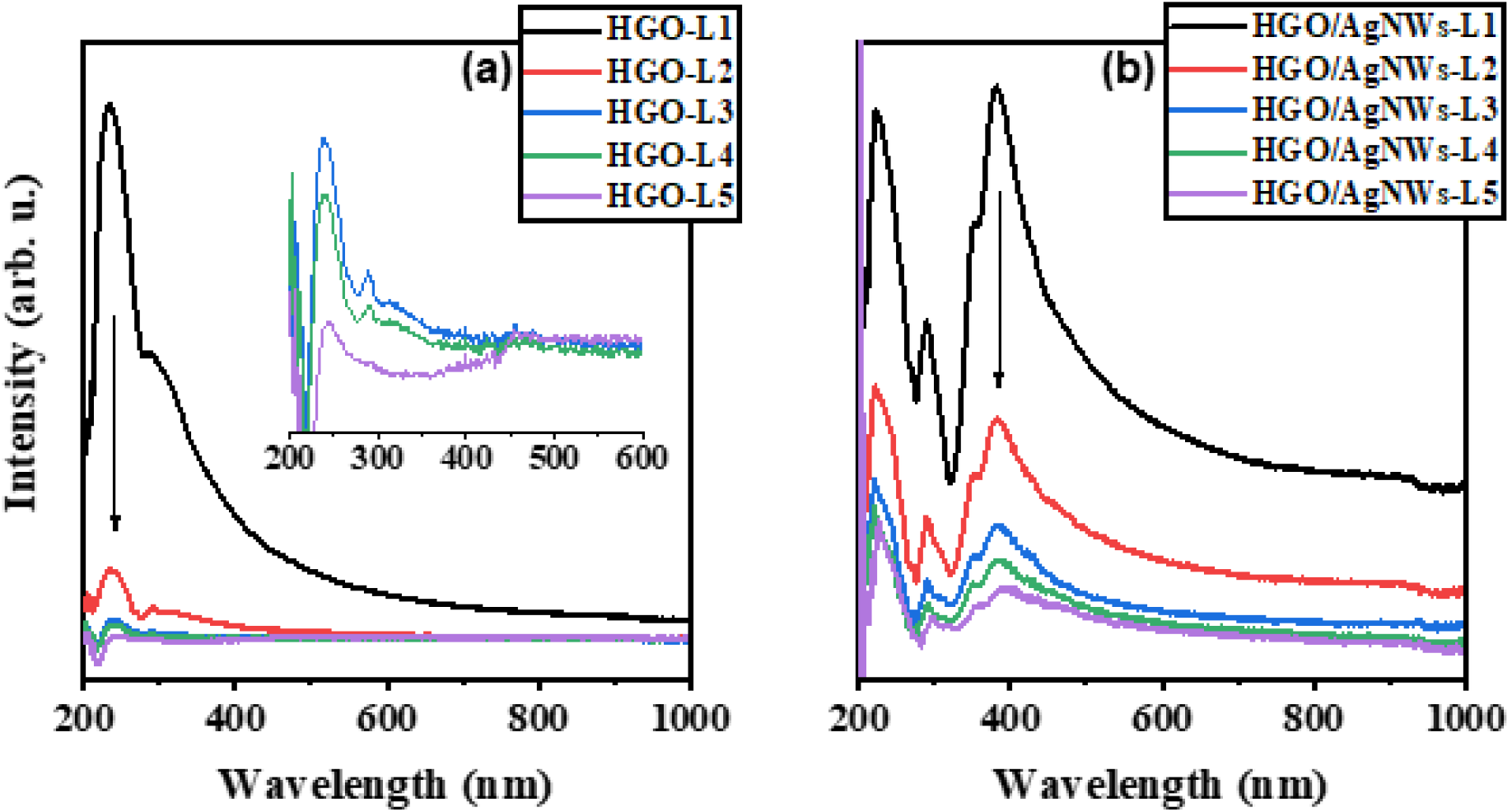
Ultraviolet-visible (UV-Vis) absorption spectra of residual wash-water dispersions collected after successive washing cycles from (a) cotton fabric (CF) treated with hydroxyl-rich graphene oxide at 3 mg/mL (CF-HGO3) and (b) cotton fabric (CF) treated with hydroxyl-rich graphene oxide at 3 mg/mL followed by silver nanowires (AgNWs) deposition (CF-HGO3/AgNWs). The spectra indicate rapid removal of excess hydroxyl-rich graphene oxide (HGO) in the first cycle and persistence of silver nanowire (AgNWs)-related bands in the combined coating during subsequent washing cycles.

### 3.3 Surface morphology before and after washing

SEM analysis corroborated the optical evidence of surface functionalization. The AgNWs-treated fabric displayed a visible silver-gray appearance after treatment, suggesting successful deposition of metallic nanostructures onto the textile surface. Before washing, the SEM images showed silver nanowires distributed across the fibers, including regions with entangled wire-like structures. After the washing cycles, the larger aggregates were markedly reduced, but individual nanowires remained randomly attached to the cotton fibers. This morphological pattern is consistent with the wash-water spectra, indicating partial removal of excess AgNWs and persistence of a retained surface-associated coating.

**Figure 3.**
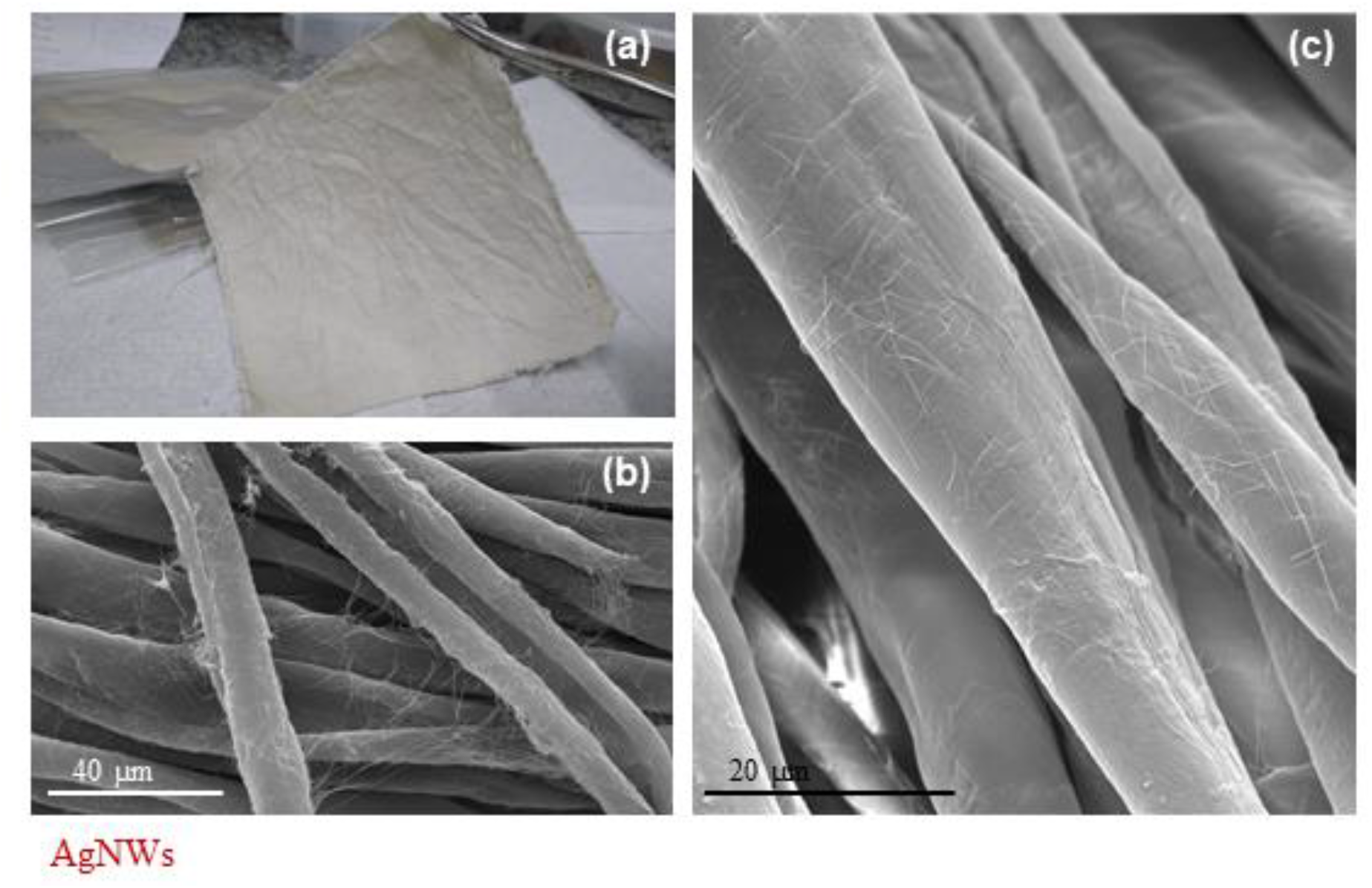
Optical and scanning electron microscopy (SEM) images of silver nanowire (AgNWs)-treated cotton fabric (CF): (a) macroscopic appearance after treatment, (b) scanning electron microscopy (SEM) image before washing, and (c) scanning electron microscopy (SEM) image after five washing cycles. The images show deposition of silver nanowires (AgNWs) on the fibers and partial retention of the coating after washing.

### 3.4 Preliminary antibacterial activity against Escherichia coli

The preliminary antibacterial assay showed a clear difference between nanomaterial-treated fabrics and the control materials retained in the final comparison. The bar plot obtained from the first experiment showed very low growth-rate values for CF-HGO3, CF-AgNWs, and CF-HGO3/Ag, whereas the PP mask and washed cotton fabric controls presented substantially higher values. Among the nanomaterial-treated samples, CF-HGO3/Ag showed the lowest measured bacterial-growth value, while CF-AgNWs also displayed near-complete inhibition. These results indicate that surface functionalization with HGO, AgNWs, or their combination can suppress E. coli proliferation under the tested conditions, with the combined treatment showing the most promising inhibitory trend in this preliminary proof-of-concept dataset.

**Figure 4.**
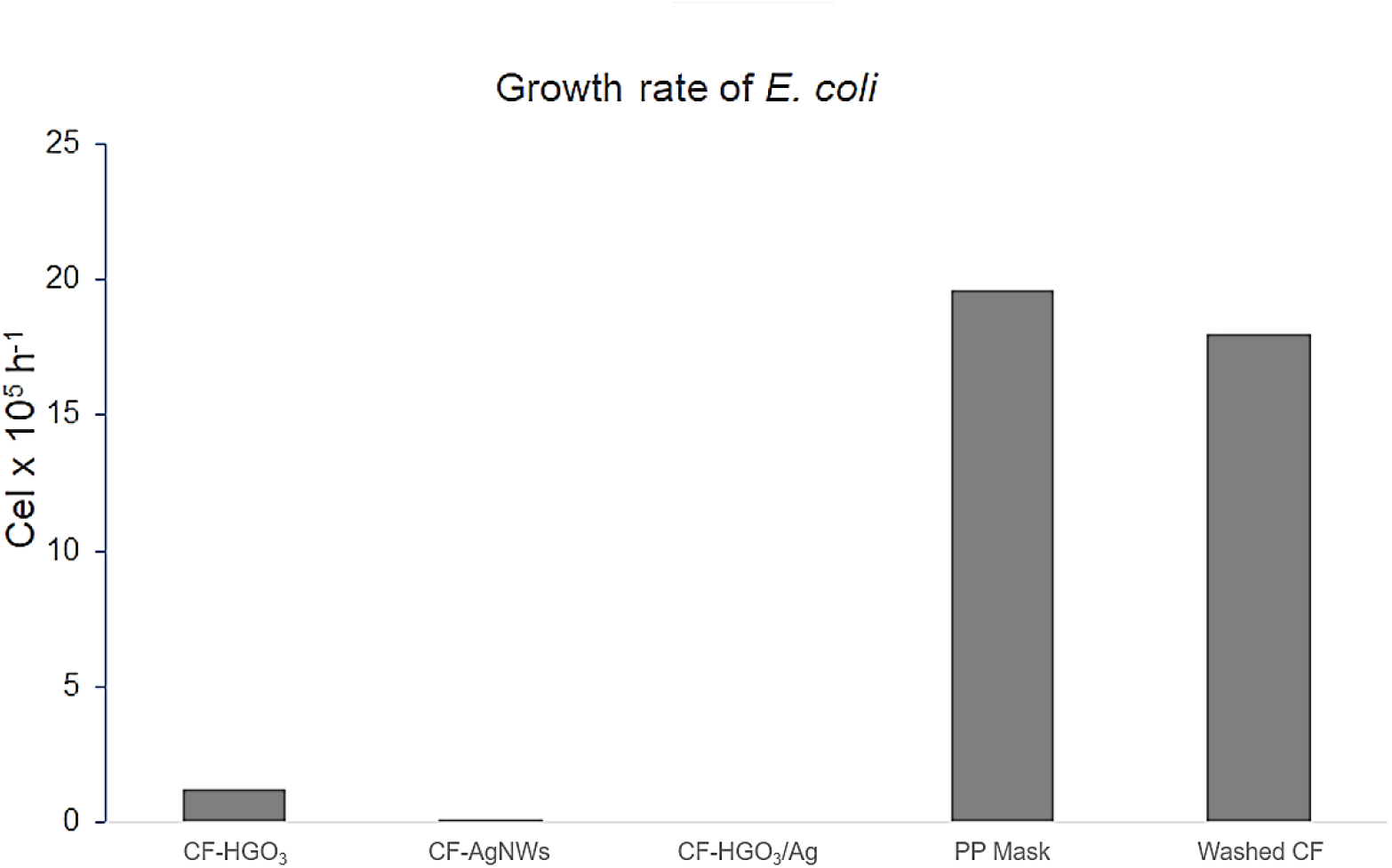
Preliminary antibacterial activity against Escherichia coli in the first retained experiment. Nanomaterial-treated fabrics, including cotton fabric (CF) treated with hydroxyl-rich graphene oxide at 3 mg/mL (CF-HGO3), cotton fabric (CF) treated with silver nanowires (AgNWs) (CF-AgNWs), and cotton fabric (CF) treated with hydroxyl-rich graphene oxide at 3 mg/mL followed by silver nanowires (AgNWs) deposition (CF-HGO3/Ag), displayed markedly lower bacterial-growth values than the polypropylene (PP) mask and washed cotton fabric (washed-CF) controls, with cotton fabric (CF) treated with hydroxyl-rich graphene oxide at 3 mg/mL followed by silver nanowires (AgNWs) deposition (CF-HGO3/Ag) showing the lowest measured value in this dataset.

### 3.5 Possible mechanisms and practical implications

The antibacterial activity observed for the treated fabrics is likely associated with complementary actions of graphene-based materials and silver nanostructures. Silver-containing nanomaterials can act through ion release, oxidative stress, and membrane damage, whereas graphene oxide derivatives may contribute to microbial inhibition through direct contact with the cell envelope, membrane perturbation, and surface-chemistry-dependent oxidative processes [2,5,8–13]. In the present study, the combined HGO/AgNWs treatment produced the lowest measured E. coli growth in the retained preliminary assay, which is compatible with a complementary contribution of both components. From an application standpoint, the present results support the feasibility of preparing cotton fabrics with partial washing resistance and measurable antibacterial activity using a simple immersion-based treatment strategy.

## 4. Conclusion

In this work, cotton fabrics were successfully functionalized with hydroxyl-rich graphene derivatives and silver nanowires through a simple immersion-based treatment approach. The combined use of UV-Vis analysis of residual wash solutions and scanning electron microscopy demonstrated that the deposited nanomaterials remained at least partially associated with the cotton fibers after repeated washing cycles, with graphene-based coatings, particularly HGO, showing stronger retention than AgNWs alone. The treated fabrics also exhibited preliminary antibacterial activity against *Escherichia coli*, indicating that the proposed surface modification strategy can impart biologically relevant functionality to the textile substrate. Among the tested conditions, the HGO/AgNWs-treated fabric showed the most promising inhibitory trend under the experimental conditions evaluated. Overall, these results provide proof of concept that cotton fabrics can be modified with hydroxyl-rich graphene derivatives and silver nanowires to yield washable materials with preliminary antibacterial performance. Further studies should address long-term durability, optimization of coating stability, broader microbiological validation, and biosafety evaluation to support future applications of these functionalized textiles.

## Notes

### Competing Interest Statement

The authors have declared no competing interest.

